# Modeling chromatin state from sequence across angiosperms using recurrent convolutional neural networks

**DOI:** 10.1101/2021.11.11.468292

**Authors:** Travis Wrightsman, Alexandre P. Marand, Peter A. Crisp, Nathan M. Springer, Edward S. Buckler

**Affiliations:** Section of Plant Breeding and Genetics, Cornell University, Ithaca, NY, USA 14853 · Funded by NSF Graduate Research Fellowship (DGE-1650441); USDA-ARS · CRediT Roles: Conceptualization, Data curation, Formal Analysis, Funding acquisition, Investigation, Methodology, Project administration, Resources, Software, Validation, Visualization, Writing - original draft, Writing - review & editing; Department of Genetics, University of Georgia, Athens, GA, USA 30602 · Funded by NSF Postdoctoral Fellowship in Biology (DBI-1905869) · CRediT Roles: Formal Analysis, Methodology, Resources, Supervision, Writing - review & editing; School of Agriculture and Food Sciences, University of Queensland, Brisbane, QLD 4072, Australia · Funded by Australian Research Council (ARC) Discovery Early Career Award (DE200101748) · CRediT Roles: Resources, Formal Analysis, Writing - review & editing; Department of Plant and Microbial Biology, University of Minnesota, Saint Paul, MN, USA 55108 · Funded by NSF IOS-1934384 · CRediT Roles: Methodology, Resources, Writing - review & editing; Section of Plant Breeding and Genetics, Cornell University, Ithaca, NY, USA 14853; Institute for Genomic Diversity, Cornell University, Ithaca, NY, USA 14853; Agricultural Research Service, United States Department of Agriculture, Ithaca, NY, USA 14853 · Funded by USDA-ARS · CRediT Roles: Conceptualization, Funding acquisition, Methodology, Supervision, Writing - review & editing

## Abstract

Accessible chromatin regions are critical components of gene regulation but modeling them directly from sequence remains challenging, especially within plants, whose mechanisms of chromatin remodeling are less understood than in animals. We trained an existing deep learning architecture, DanQ, on leaf ATAC-seq data from 12 angiosperm species to predict the chromatin accessibility of sequence windows within and across species. We also trained DanQ on DNA methylation data from 10 angiosperms, because unmethylated regions have been shown to overlap significantly with accessible chromatin regions in some plants. The across-species models have comparable or even superior performance to a model trained within species, suggesting strong conservation of chromatin mechanisms across angiosperms. Testing a maize held out model on a multi-tissue scATAC panel revealed our models are best at predicting constitutively-accessible chromatin regions, with diminishing performance as cell-type specificity increases. Using a combination of interpretation methods, we ranked JASPAR motifs by their importance to each model and saw that the TCP and AP2/ ERF transcription factor families consistently ranked highly. We embedded the top three JASPAR motifs for each model at all possible positions on both strands in our sequence window and observed position- and strand-specific patterns in their importance to the model. With our cross-species “a2z” model it is now feasible to predict the chromatin accessibility and methylation landscape of any angiosperm genome.

## Introduction

Accessible chromatin regions (ACRs) are known to play a critical role in eukaryotic gene regulation but their comprehensive identification in plants remains a challenge [<u>1,2</u>]. Current methods to assay chromatin accessibility are highly environment-specific and relatively expensive compared to DNA sequencing, limiting the number of species or conditions that can be investigated. Assaying chromatin accessibility in plants comes with additional unique challenges: the cell wall makes plant nuclei hard to isolate and many active transposon families shuffle, create, and destroy regulatory regions over time [<u>3</u>]. Regions that lack DNA methylation are known to be stable over developmental time and overlap significantly with ACRs in plants with larger genomes [<u>4</u>], suggesting they may contain a superset of ACRs across cell-types. Computational models capable of predicting chromatin accessibility and methylation state directly from DNA sequence would enable a wide range of previously-intractable studies on gene regulation across evolutionary time as well as estimation of non-coding variant effects for use in contexts such as breeding. Plants also provide an excellent system to study the genetic basis of adaptation [<u>5</u>]. Now that it is feasible to assemble genomes of thousands of species, regulatory regions that control adaptation can be identified, providing valuable insight on how to breed crops resilient to climate change. Recent advances in machine learning, particularly deep learning, have catalyzed a vast number of applications to biological prediction, including mRNA abundance [<u>6,7,8</u>], chromatin state [<u>9,10,11</u>], and transcription factor (TF) binding [<u>12</u>] directly from DNA sequence. Many of these models have so far only been trained within a single species to predict within the same species, usually utilizing held-out chromosomes as a test set to control for sequence relatedness.

At a high level, plant chromatin has characteristics similar to animal chromatin: chromatin is organized into hierarchical compartments, distal regulatory regions are colocalized to genes through chromatin looping, and various histone modifications signal a wide variety of local chromatin states. However, the exact mechanisms driving chromatin accessibility are known to be quite different in terms of specific histone modifications [<u>13</u>], pioneer factors [<u>14</u>], and chromatin looping mechanisms [<u>15</u>]. Because of these differences, plant-specific chromatin accessibility models are likely to be necessary.

We know that transcription factor binding sites are strongly conserved across evolutionary time [<u>16</u>, <u>17</u>] and highly enriched in ACRs [<u>18</u>]. Certain deep learning model architectures, such as convolutional neural networks (CNN), have already been shown effective for predicting chromatin accessibility within species by recognizing important motifs [<u>9,10</u>] and their spatial relationships [<u>19</u>]. Previous work [<u>17,20</u>] has observed that CNNs require much larger training data sets than earlier model architectures to achieve equivalent or better performance. By incorporating multiple species into the training data we not only increase the number of observations but also the total evolutionary time between observations, which reduces confounding neutral variation within conserved sequences. For the purposes of predicting regulatory regions in unobserved plant species, training a model across species will be critical to learn important motifs and syntax that are conserved across longer evolutionary time periods. Therefore, we predicted that previously-published deep learning architectures could work well across species and make accurate chromatin accessibility and methylation predictions in related unobserved species. DanQ [<u>10</u>] is a recurrent CNN that has already been shown to be able to more accurately predict a number of genomic labels, including chromatin accessibility and DNA methylation, in the human genome than standard CNNs like DeepSEA [<u>9</u>].

Here, we train DanQ to predict chromatin accessibility using leaf ATAC-seq data from 12 angiosperm species [<u>13</u>], comparing the performance of within-species models to across-species models. We also train DanQ to predict unmethylated regions using methylation data from 10 angiosperm species, including 5 previously-published grasses [<u>4</u>]. Using a maize single-cell ATAC (scATAC) accessibility atlas [<u>21</u>], we see that the accessibility model has similar performance across cell-types but is highly variable across regions with different levels of cell-type specificity. Using various interpretation methods designed for CNNs, we compare and contrast which motifs were important across angiosperms for predicting chromatin accessibility in leaves or methylation state. Our pan-angiosperm chromatin state models are an important stepping stone towards a better understanding of gene regulation and adaptation.

## Results

### Recurrent convolutional neural networks accurately model chromatin state across species

To train a successful chromatin state classifier, we needed to choose a window size that balanced genomic context with resolution. We tested a few different model configurations and decided upon 600 basepair windows because higher window sizes showed diminishing returns on performance while decreasing our effective resolution (Figure <u>S1</u>). We preprocessed the ATAC-Seq and unmethylated peaks by taking the midpoint and symmetrically extending to half the window size in both directions to obtain our postive observations. Negatives were sampled from the rest of the genome. After preprocessing we had 26,280 training regions per species (315,360 total) for the cross-species accessibility models and 35,652 training regions per species (356,520 total) for the methylation models, split evenly between classes.

As a baseline for comparison to previous, within-species, chromatin state CNN models as well as our across-species models, we trained within-species DanQ model configurations for each of the angiosperm species in our data. We also trained across-species model configurations each using a different species as a test set. Generally, we observed that a given across-species model has a comparable, if not superior, area under the precision-recall curve (auPR) to the within-species model (Figure <u>1</u>, top middle and top right). While auPR across species varies substantially, they are also within the range of those observed in the original DanQ and DeepSEA human models and superior to the bag-of-kmers model in *Zea mays* (Figure <u>S3</u>). We also see that both within-species and across-species performance decreases as genome size increases (Figure <u>S4</u>). When comparing the accessibility and hypomethylation models, we see the same trends in performance for each species.

**Figure 1:**
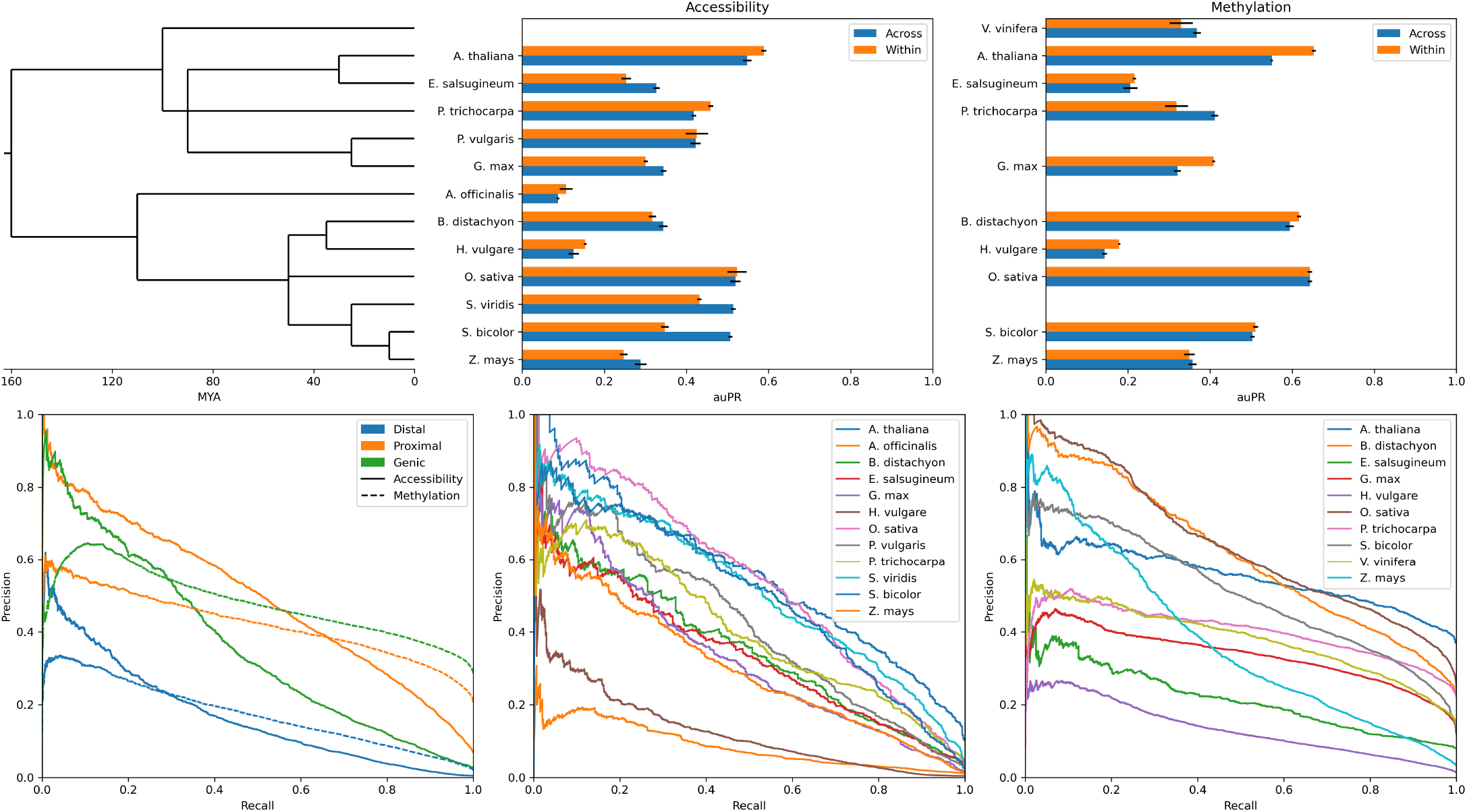
Performance of the cross-species chromatin state classifiers. The top middle and top right show the mean and standard error (due to variability in the stochastic model parameter initialization processes) of the auPR for the accessibility and methylation models, respectively, per species for both the within- and across-species training configurations. The bottom left is the precision-recall curve across all hold-out species for the across-species models, split by distance class and chromatin feature. The bottom middle and bottom right are the precision-recall curves for the across-species accessibility and methylation models, respectively, split by species.

To see if the models were more accurate in predicting accessible or unmethylated regions near or within genes, where these regions are known to be enriched, we looked at the precision-recall curves across different distance classes (genic, proximal, or distal). Observations were labeled as genic if more than half of the range overlapped with a gene annotation, as proximal if not genic and more than half of the range was within the proximal cutoff (2kb), and as distal if neither genic nor proximal. We see that the across-species models for both chromatin features perform the worst on distal regions, but show contrasting results on the genic and proximal regions (Figure <u>1</u>, bottom left). This could be driven by the imbalanced distribution of regions between the distance classes, with accessible regions biased towards the proximal class and unmethylated regions towards the genic class (Figure <u>S5</u>). In particular, *Hordeum vulgare* has proportionally many more distal accessible and unmethylated regions, which could explain the lower overall performance. The across-species accessibility models are very precise when calling inaccessible chromatin, with most of the errors being false-positives, particularly in distal regions (Figure <u>S6</u>). We see a much different result in the methylation model, which shows only a slight bias towards false positives.

To control for potential *trans*-driven transposon silencing, we tested a two-step model that takes the predictions of the a2z model and then masks them with zeros if they overlap annotated transposons in *Z. mays*. We see that these two-step repeat-masked models do much better (ΔauPR 0.15 for accessibility and 0.07 for methylation) than the naive models (Figure <u>S7</u>), suggesting a relatively straightforward way to reduce false positives in larger plant genomes with more transposon-derived sequence.

Finally, we wanted to assess how far out in evolutionary time the angiosperm model could work. We ran the model against ATAC-Seq data from *Saccharomyces cerevisiae* and a *Homo sapiens* GM12878 cell line [<u>22</u>]. We see the plant-trained model has some ability (Figure <u>S8</u>) to predict chromatin accessibility in yeast (auPR 0.21), if not human cell-lines (auPR 0.02).

### Leaf-trained models struggle to predict cell type-specific accessible chromatin regions

Knowing the a2z models are capable of working across species, we then asked how well the leaf-trained accessibility models could work across cell types. We used scATAC-Seq data from six maize organs [<u>21</u>] as a multi-cell type test set for our single-tissue model. Using a model trained on every species with ATAC-seq data except *Z. mays*, we predicted the accessibility of each scATAC peak as well as negatives sampled from the rest of the genome. Looking at the area under the threshold-recall curve we see that the model does better on peaks that are accessible across many cell types, with a sharp decrease in peaks only accessible in five or fewer cell types, which are likely to be a mix of false positives and highly cell type-specific peaks (Figure <u>2</u>, left). The model does best on peaks that are generally open across many cell types, which comprise a largest portion of the training data (Figure <u>S9</u>). This is clearly shown when looking at the overall precision-recall curves in the best (guard cell) and worst (trichoblast) cell types, as well as a union of all cell types. There is not a substantial difference between the three (Figure <u>2</u>, right).

**Figure 2:**
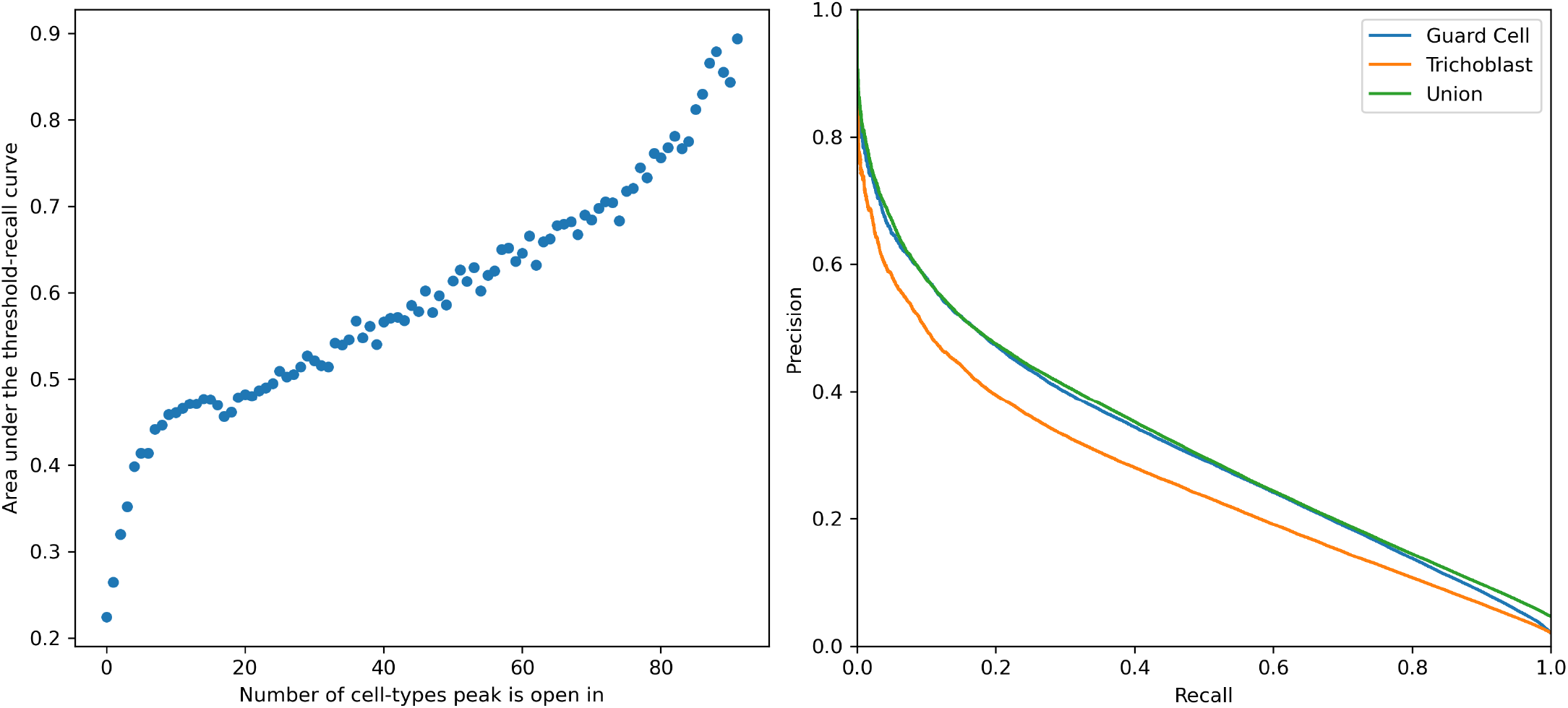
Cross-cell type performance of the *Zea mays* accessibility model. The left plot shows the area under the threshold-recall curve for each set of peaks grouped by the number of cell types they are accessible in. The right plot shows the precision-recall curves for peaks accessible in the guard cell (best) and trichoblast (worst) cell types, as well as peaks open in any cell type (union).

### Interpretation methods reveal important conserved and species-specific motifs

Although chromatin state models that work across angiosperms are a useful tool, we may be able to gain new insights into chromatin biology by dissecting what motifs and higher-order motif patterns the model is learning to use to separate accessible from inaccessible chromatin or unmethylated from methylated regions. We started with the attribution tool TF-MoDISco to identify important motifs in the *Z. mays* and *Arabidopsis thaliana* test sets using their respective held-out models. While TF-MoDISco qualitatively identified many important motifs (Figure <u>S10</u>), most of them ranked similarly by attribution score and therefore could not be quantitatively compared in terms of effect size or importance relative to each other.

To obtain better estimates of sequence effect size, we developed a method that masks sliding windows across a set of sequences and evaluates the change in the model prediction, which we refer to as the kmer occlusion method. Using a kmer size of 10bp, representing a common estimate of core binding site length, we ran a kmer occlusion to get effect sizes for each kmer in the test set, binned kmers into “high-effect” and “null-effect”, and then scanned them for matches to JASPAR 2020 CORE *plantae* [<u>23</u>] binding motifs. For our accessibility models, we see that about 20-40% of high-effect kmers match with JASPAR motifs while our methylation models generally seem to have poor matching between JASPAR motifs and high-effect kmers (Figure <u>S12</u>). To look at how similar the high-effect kmers were between chromatin features and species, we used k-medoids to get a subset of representative kmers and then visualized the distances between them using multidimensional scaling. Surprisingly, the high-effect kmers across species and chromatin features cluster together, with slight separation between methylation and accessibility (Figure <u>3</u>, left). However, there is no separation between species (Figure <u>S11</u>) nor monocots and dicots (Figure <u>3</u>, middle/right) for either chromatin feature.

**Figure 3:**
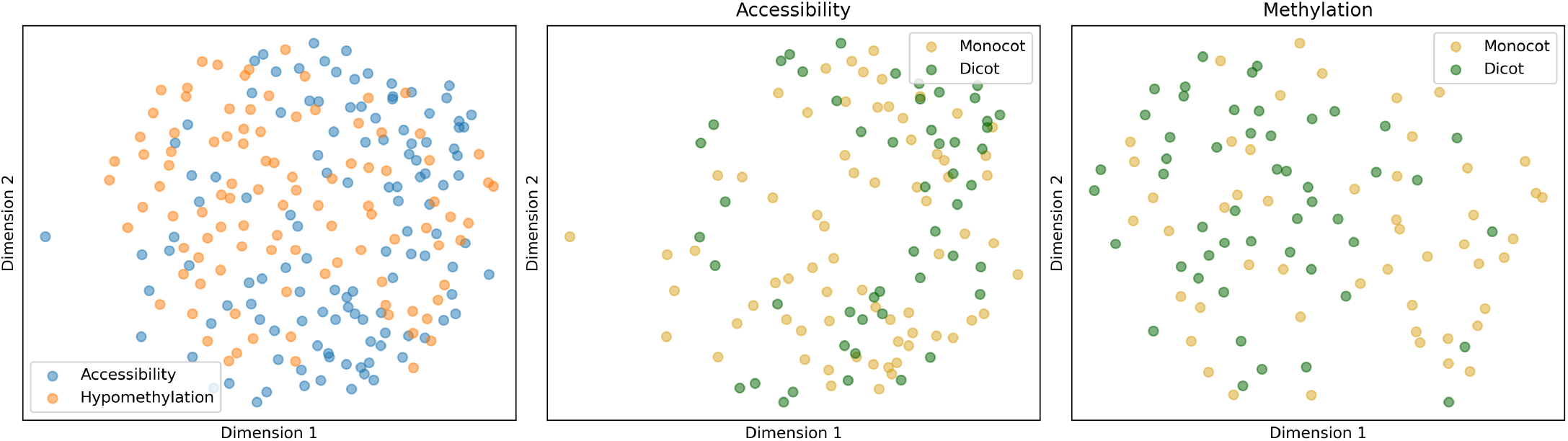
Multidimensional scaling of the high-effect medoid kmer distance matrix across all species and chromatin feature model combinations. Each point is a high-effect kmer in a given species and chromatin feature combination.

To understand which known biological motifs were being recognized as important to the model, we used a recently-developed model interpretation method known as Global Importance Analysis (GIA) [<u>2</u> <u>4</u>]. First, we ranked JASPAR motifs by their max global importance across all positions for each model (Table <u>1</u>) and see both species-specific and common TFs across the models. One of the most remarkable observations is that the top 10 motifs in the *A. thaliana* model are all from the TCP family. The *Z. mays* accessibility model also ranked TCP motifs in the top 10 but behind Dof-type motifs. The *A. thaliana* and *Z. mays* methylation models rank the same two motifs at the top and share mostly the same families between the rest. Next, we looked at the positional effects of the top three TFs across *A. thaliana* accessibility (Figure <u>4</u>, top left) and methylation (bottom left) as well as *Z. mays* accessibility (top right) and methylation (bottom right). The most striking feature is the sawtooth pattern seen across both species and chromatin feature models, however the cause of this pattern is unclear. The *A. thaliana* accessibility model shows a clear bias towards the center of the accessible regions for the top three TFs while the other models are not as consistent.

**Table 1:** Top 10 JASPAR motifs for four pan-angiosperm models ranked by max global importance across all possible embedding positions. TF family or class (if family was not available) according to JASPAR is shown in parentheses under each TF.

| Rank | Accessibility<br>*A. thaliana* | Accessibility<br>*Z. mays* | Methylation<br>*A. thaliana* | Methylation<br>*Z. mays* |
| --- | --- | --- | --- | --- |
| 1 | TCP1<br>(TCP) | AT5G66940<br>(Dof-type) | ERF104<br>(AP2/ERF) | ERF104<br>(AP2/ERF) |
| 2 | TCP14<br>(TCP) | OBP3<br>(Dof-type) | AT4G18450<br>(AP2/ERF) | AT4G18450<br>(AP2/ERF) |
| 3 | At1g72010<br>(TCP) | AT1G69570<br>(Dof-type) | ERF9<br>(AP2/ERF) | RAP211<br>(AP2/ERF) |
| 4 | TCP21<br>(TCP) | OBP1<br>(Dof-type) | BPC5<br>(BBR-BPC) | BPC5<br>(BBR-BPC) |
| 5 | TCP19<br>(TCP) | AT2G28810<br>(Dof-type) | ERF2<br>(AP2/ERF) | ERF9<br>(AP2/ERF) |
| 6 | TCP7<br>(TCP) | AT5G02460<br>(Dof-type) | LEP<br>(AP2/ERF) | ESE1<br>(AP2/ERF) |
| 7 | At2g45680<br>(TCP) | TCP1<br>(TCP) | BPC1<br>(BBR-BPC) | AT5G66940<br>(Dof-type) |
| 8 | TCP20<br>(TCP) | At1g72010<br>(TCP) | ESE1<br>(AP2/ERF) | BPC1<br>(BBR-BPC) |
| 9 | OJ1581_H09.2<br>(TCP) | TCP21<br>(TCP) | ERF10<br>(AP2/ERF) | ERF2<br>(AP2/ERF) |
| 10 | TCP2<br>(TCP) | BPC5<br>(BBR-BPC) | BPC6<br>(BBR-BPC) | LEP<br>(AP2/ERF) |

**Figure 4:**
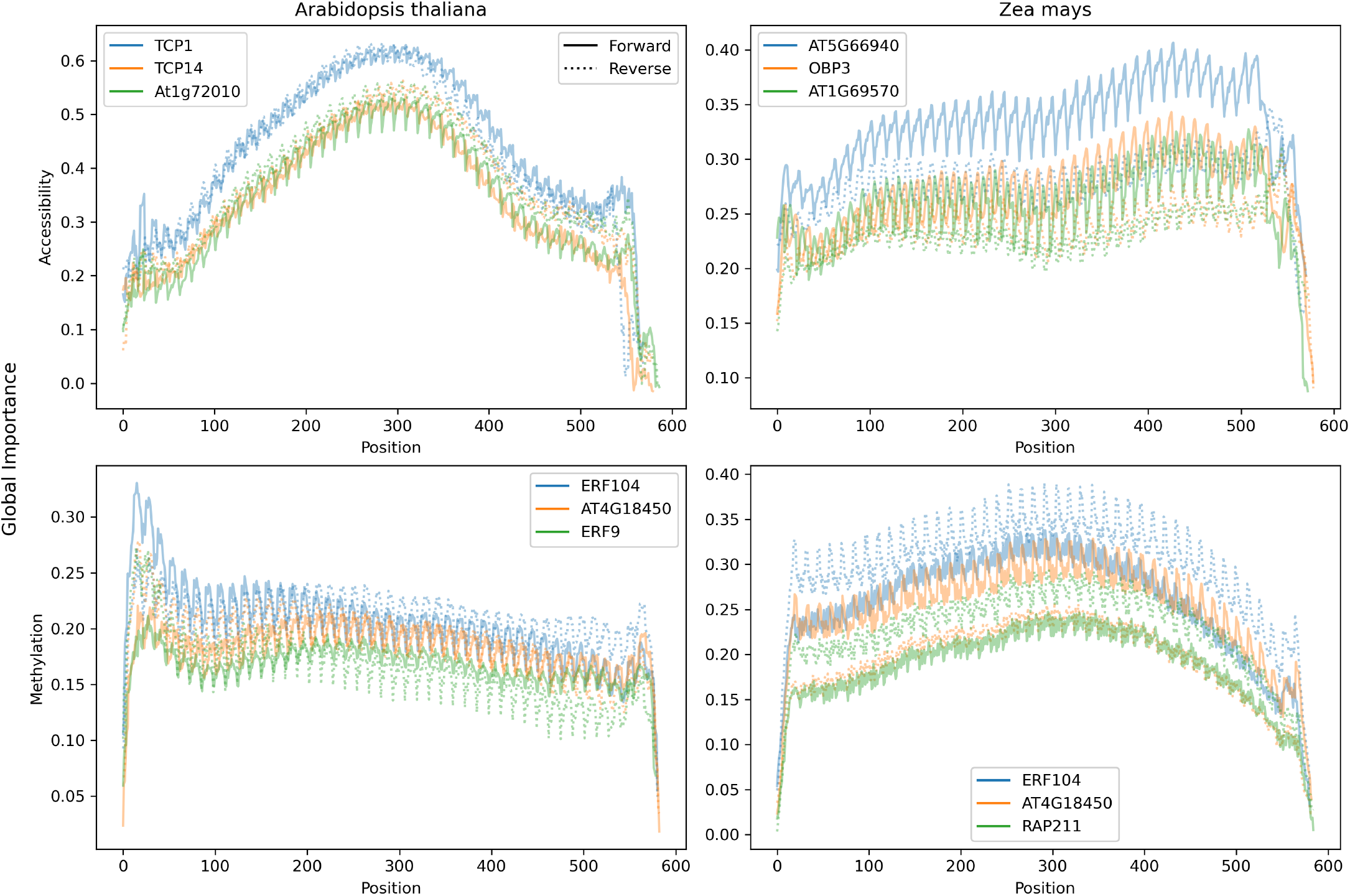
Positional Global Importance Analysis plots for *A. thaliana* (left) and *Z. mays* (right) accessibility (top) and methylation (bottom). The solid and dotted lines represent the importance scores for the positive and negative strand, respectively. Only the top three JASPAR motifs ranked by the maximum global importance across the sequence were plotted.

## Discussion

We have shown that recurrent CNNs, DanQ in particular, are an effective architecture on which to base cross-species sequence to chromatin state models. By incorporating sequence data from multiple species we not only increase the size of our training data set, a critical factor for deep learning models, but also reduce the amount of confounding neutral variation around functional motifs. Being able to predict chromatin state across species also opens the door for studies of regulatory regions in additional angiosperm species with only genomic sequence data. Beyond angiosperms, the a2z model’s predictive ability in yeast suggests it is capable of working effectively across wide evolutionary timescales. Unsurprisingly, we noticed that the performance across different peak classes relates to their relative abundance in the training set. Future work looking at ways to balance or weight observations in rarer peak classes would likely improve the generalizability of the models. This is particularly important for working towards better cross-tissue chromatin state models, where the tissue-specific peaks are usually the minority in any given data set, as well as with larger genomes, where distal peaks are more prevalent.

Further, most sequence-based model architectures, including DanQ, only take in *cis* sequence, which is known [<u>25</u>] to account for only a portion of the variation in local chromatin state. Model architectures that can effectively incorporate *trans* factors, such as chromatin-remodeling TFs on neighboring regulatory elements [<u>26</u>] or small RNA silencing [<u>27</u>], will likely surpass current methods but their cross-species applicability remains an open question. By far the most prevalent error of the accessibility models in particular is calling false-positives, which may be due to lack of *trans* information. A portion of these false-positives may also be undercalled ATAC-Seq peaks that are open in very specific cell-types, since the peaks from Lu *et al.* 2019 were called with relatively conservative thresholds.

Interpreting deep learning models remains a challenge, but is an especially critical one to overcome. Here we use occlusion and perturbation-based methods instead of gradient-based approaches like TF-MoDISco and saliency maps to trade longer computational times for reduced noise [<u>28</u>] in effect estimates. Particularly since eukaryotic TF binding sites are known to be degenerate [<u>29</u>], SNP effect sizes in regulatory sequences are likely to be small and harder to estimate accurately with our limited data. The lack of separation between clades and species in the MDS plots for each chromatin feature is not too surprising. The cross-species models must learn to prioritize motifs that are generalizable across species and so potential species- or clade-specific motifs are ignored. The sawtooth pattern, which is stronger in some TFs than in others, could be a manifestation of the model learning a helical face bias for specific TF binding. Further controls will be necessary to investigate that hypothesis, as the pattern may also be an artifact of the max pooling or LSTM layers. Not all of the pGIA results agree with current theory. For example, some of the motifs have a noticeable strand bias, but enhancers are known to operate in an orientation-independent [<u>30</u>] manner. Given some of them are relatively simple motifs, it is possible that these matches are surrogates for important non-binding motifs. We chose to rank JASPAR motifs by maximum global importance across the sequence as a rough estimate for importance to regulating the given chromatin feature state, though other methods of ranking could be preferrable depending on the use case. Since positive observations are created by extending from the midpoint, the effect of TFs that bind to the center of accessible or unmethylation regions will be easier to estimate because they are more aligned across the test set sequences. In contrast, TFs that bind to the edges of accessible or unmethylation regions are not aligned since the lengths of the true, unextended ATAC-Seq peaks are not equal.

The top 10 JASPAR motifs are very different between the features but remarkably similar between the species within each feature. Of the two known [<u>31,32,33</u>] plant pioneer transcription factors (LEC1 and LEAFY), only LEAFY is present in JASPAR, but does not show up in the top 10 motifs for any of the models. This is not unexpected as it is a floral TF and our models are trained on leaf accessible regions. The strong presence of the TCP family in the highly ranked accessibility TFs is promising, since they are known [<u>34</u>] to be involved in chromatin remodeling. What role the Dof-type TFs play in accessibility is still unclear due to the wide variety of roles they play [<u>35</u>]. The shared top two motifs between the methylation models have evidence that they are involved in plant pathogen response [<u>36,37</u>]. Knowing that plant immunity genes are among the most variable [<u>38</u>], it would be interesting to see if these unmethylated regions are harboring a large library of rapidly inducible resistance genes that remain mostly inaccessible until needed. With the high similarity in binding motifs by definition within families, it is quite possible that some highly ranked TFs are false positives due to association with the few causal TFs in the same family. While it is useful to use JASPAR motifs as specific testable hypotheses, there are only 530 motifs in the database and with the lowest estimates of angiosperm TF gene count starting at about 1,500 [<u>39</u>], critical TFs may still be missing.

Moving forward, more focus is necessary on collecting high-quality accessible regions across a variety of cell-types to train models that are capable of generalizing across tissues as well as species. With the release of highly-accurate protein-folding models such as AlphaFold2 [<u>40</u>], the missing species-specific TF binding motifs in any genome may finally be feasible to estimate using simulated DNA docking approaches. Now that many deep learning-based approaches borrowed from other fields [<u>12,41</u>] have been shown to be successful in mapping genomic sequence to a variety of cellular phenotypes, better interpretation methods to assess what these black box models are learning will be important to optimize towards more biologically-relevant architectures.

## Materials and Methods

### Software environment

The software environment for the experiments was managed by <u>conda</u> (v4.10.3). Packages were downloaded from the conda-forge [<u>42</u>] and bioconda [<u>43</u>] channels. Software versions not explicitly mentioned in the methods are defined in the conda environment files in the companion code repository on <u>Zenodo</u>.

### Raw data

The angiosperm ATAC-seq peaks [<u>13</u>] were downloaded from NCBI GEO accession GSE128434. Genomes and annotations for *Arabidopsis thaliana* (TAIR10) [<u>44</u>], *Eutrema salsugineum* (v1.0) [<u>45</u>], *Phaseolus vulgaris* (v1.0) [<u>46</u>], *Glycine max* (Wm82.a2.v1) [<u>47</u>], *Brachypodium distachyon* (v3.0) [<u>48</u>], *Oryza sativa* (v7.0) [<u>49</u>], *Setaria viridis* (v1.0) [<u>50</u>], *Populus trichocarpa* (v3.0) [<u>51</u>], and *Sorghum bicolor* (v3.1 and v3.1.1) [<u>52</u>] were downloaded from Phytozome. Reference genomes and annotations for *Zea mays* (AGPv4.38) [<u>53</u>] and *Hordeum vulgare* (IBSC_v2) [<u>54</u>] were downloaded from Ensembl Plants. The genome and annotation for *Asparagus officinalis* (v1.1) [<u>55</u>] was downloaded from the Asparagus Genome Project website. Unmethylated regions (UMRs) for the grasses were downloaded from the supplemental information of Crisp *et al.* 2020 [<u>4</u>]. For the unmethylated regions, the *Z. mays* AGPv4 genome and annotation was downloaded from MaizeGDB. The *Vitis vinifera* genome and annotation (Genoscope.12X) [<u>56</u>] were downloaded from the Genoscope website.

JASPAR 2020 Core *Plantae* [<u>23</u>] motifs and clusters were downloaded from the JASPAR website. Maize AGPv4 RepeatMasker annotations were downloaded from NCBI. Yeast and human cell-line GM12878 ATAC-seq peaks [<u>22</u>] were downloaded from NCBI GEO accession GSE66386. The yeast (sacCer3 April 2011) [<u>57</u>] and human (hg19) [<u>58</u>] genomes were downloaded from NCBI. Maize scATAC-seq peaks [<u>21</u>] were downloaded from NCBI GEO accession GSE155178. Genome files were indexed using samtools [<u>59</u>].

### UMR calling on non-grass species

UMR analysis on the non-grass species was performed as per Crisp *et al.* 2020. Briefly, sequencing reads were trimmed and quality checked using Trim galore! (0.6.4_dev), powered by cutadapt (v1.18) [<u>60</u>] and fastqc (v0.11.4). For all libraries, 20bp were trimmed from the 5’ ends of both R1 and R2 reads and aligned with bsmap (v2.74) [<u>61</u>] to the respective genomes with the following parameters: - v 5 to allow 5 mismatches, −r 0 to report only unique mapping pairs, and −p 1 and −q 20 to allow quality trimming to Q20. Output SAM files were parsed with SAMtools [<u>62</u>] fixsam, sorted, and then indexed. Picard MarkDuplicates [<u>63</u>] was used to remove duplicates, <u>BamTools</u> filter to remove improperly paired reads, and bamUtil clipOverlap [<u>64</u>] to trim overlapping reads so as to only count cytosines once per sequenced molecule in a pair for PE reads. The methylratio.py script from bsmap was used to extract per-site methylation data summaries for each context (CH/CHG/CHH) and reads were summarised into non-overlapping 100bp windows tiling the genome. WGBS pipelines are available on <u>GitHub</u>. To identify unmethylated regions, each 100bp tile of the genome was classified into one of six domains or types: “missing data” (including “no data” and “no sites”), “High CHH/RdDM”, “Heterochromatin”, “CG only”, “Unmethylated” or “intermediate”, in preferential order as per Crisp *et al.* 2020 [<u>4</u>].

### Training data preprocessing

Interval manipulation was done using a combination of the GNU coreutils, gawk, and bedtools [<u>65</u>]. We created our positive observations by symmetrically extending each accessible or unmethylated region from the midpoint by half of the window size (300, 600, or 1000 bp). Our negative observations are randomly sampled from the rest of the genome not covered by the union of the resized positive observations and the original peaks. Observations were labeled as genic if more than half of the range overlapped with a gene annotation, as proximal if not genic and more than half of the range was within the proximal cutoff (2kb), and as distal if neither genic nor proximal. Previous work [<u>20,66</u>] has shown that classifiers train best on balanced sets with an equal number of positive and negative examples, but should be tested on the true class distribution to get an accurate performance estimate. Therefore, for the across-species models, we randomly sampled 6% of the observations and divided them equally between a validation and test set. For the within-species models we randomly chose a hold-out chromosome to follow best practice for reducing contamination of related sequences between the training and test sets. As a heuristic to select held-out chromosomes across genome assemblies of varying contiguity, we randomly select within chromosomes that are at least a million basepairs long and have more than five positive observations. We then downsampled the remaining observations to obtain a training set for the across-species models with a balanced representation of species and target class. Ns were encoded as vectors with equal probability assigned to each base as opposed to all zeros, which is another common practice. Sequences were extracted using BioPython [<u>67</u>] and pyfaidx [<u>68</u>]

### Training and evaluating the DanQ architecture

The DanQ architecture was implemented using the keras [<u>69</u>] API of tensorflow [<u>70</u>]. The across-species models were tested on a given species and trained on the remainder. Within-species models were tested on a held-out chromosome and trained on the other chromosomes. Since our ratio of accessible to inaccessible chromatin observations is heavily unbalanced, we focus more on the area under the precision-recall curve (auPR) to measure model performance as opposed to the more commonly-reported area under the receiver operating characteristic curve (auROC). Performance metrics were measured using scikit-learn [<u>71</u>] and curves were plotted using matplotlib [<u>72</u>]. Each model was trained three times to obtain an estimate of variability in performance due to the stochastic nature of the model variable initialization. For comparison between models we used the first of the three trained models.

The bag-of-kmers model was trained and tested independently on the within-species *Z. mays* accessibility and methylation training data using code adapted from Tu *et al.* 2020 [<u>12</u>] and compared to the within-species *Z. mays* accessibility and methylation models. For the two-step masked model comparison we masked the *Z. mays*-held-out accessibility and methylation model predictions to zero if more than half of a region overlapped with an annotated repeat from RepeatMasker. We used pybedtools [<u>73</u>] to compute overlaps between the test set and the repeats. We preprocessed the yeast and human cell-line ATAC-seq peaks in the same manner as the angiosperm ATAC-seq peaks and used the *Z. mays*-held-out model to make predictions on the yeast and human peaks.

The grasses accessibility model was trained and evaluated in the same manner as the across-species angiosperm accessibility model but restricted to only grass species. The “balDist” accessibility model extended the training data balancing to distance class in addition to chromatin state, meaning the training data had equal representation for each species, distance class (genic, proximal, distal), and target class (accessibile/inaccessible or unmethylated/methylated). The “exp” accessibility model changed the activation function on the convolutional layer from ReLU to exponential. The “all_v_AtZm” accessibility model was tested on *A. thaliana* and *Z. mays* and trained on the rest of the angiosperm species.

The dendrogram in Figure <u>1</u> was plotted using the Phylo package of Biopython [<u>74</u>].

### Analysis of maize scATAC-Seq data

scATAC-seq peaks were preprocessed in the same manner as the other peaks to generate uniform 600 bp regions. Peaks were classified as open in a cell-type if their CPM (counts per million, a normalized depth measurement) value was greater than in that cell-type, which would represent no reads observed in that peak in that cell-type, based on the methods reported in Marand *et al.* 2021 [<u>21</u>]. Accessibility was predicted using the *Z. mays*-held-out model.

### TF-MoDISco and kmer occlusion

We ran TF-MoDISco [<u>75</u>] with a sliding window size of 15bp, a flank size of 5bp, and a target seqlet FDR of 0.15. For converting seqlets to patterns, we set “trim_to_window_size” to 15bp, “initial_flank_to_add” to 5bp, and specified a final minimum cluster size of 60.

The kmer-occlusion method involves masking (replacing with N’s) a sliding kmer across each sequence in a given model’s test set. The difference between the model’s masked and unmasked prediction is the kmer’s “effect size”. We ran the kmer-occlusion method with a kmer size of 10bp on all species and chromatin feature pairs. The top 5% accessibility- or methylation-reducing kmers per species and chromatin feature were classified as “high-effect” kmers. We performed an all-by-all global alignment of the high-effect kmers per species and chromatin feature using Biopython’s pairwise aligner [<u>67</u>]. Using the alignment distance matrix, we clustered these high-effect kmers into 100 representative kmers using k-medoids [<u>76</u>]. We took the 100 medoid kmers for each species and chromatin feature pair and did another all-by-all global alignment to create another distance matrix. The embedded kmer coordinates were created using the MDS function in scikit-learn’s manifold package. High-effect kmers were matched to JASPAR 2020 CORE *plantae* motifs using FIMO [<u>77</u>] and a q-value threshold of 0.05.

### Positional Global Importance Analysis

Global importance analysis (GIA) [<u>24</u>] measures the average difference in model predictions from a sampled background set of sequences to the same set with the sequence embedded within them. We ran a positional GIA (pGIA) analysis for each species and chromatin feature pair by embedding the consensus motifs of the 530 JASPAR 2020 CORE *plantae* TFs in both orientations at each possible position within 1,000 generated 600bp sequences. The 600bp sequences were generated using a profile model, where bases were sampled at each position according to their relative frequency in the model’s test set at that position. GNU parallel [<u>78</u>] was used to speed up the pGIA analysis.

JASPAR motifs were ranked by their maximum global importance across all positions. TF families and classes were obtained from the JASPAR API (v1).

### Manuscript

This manuscript was formatted with Manubot [<u>79</u>].

## Acknowledgements

This work was funded by an NSF Graduate Research Fellowship (DGE-1650441) and the USDA-ARS to T.W., an NSF Postdoctoral Fellowship in Biology (DBI-1905869) to A.P.M., an Australian Research Council (ARC) Discovery Early Career Award (DE200101748) to P.A.C., the NSF IOS-1934384 to N.M.S., and the USDA-ARS to E.S.B. The Texas Advanced Computing Center supported a portion of the compute time for the analyses with their Frontera system. Peter Koo contributed helpful comments during the analysis.

## Supplementary Information

**Figure S1:**
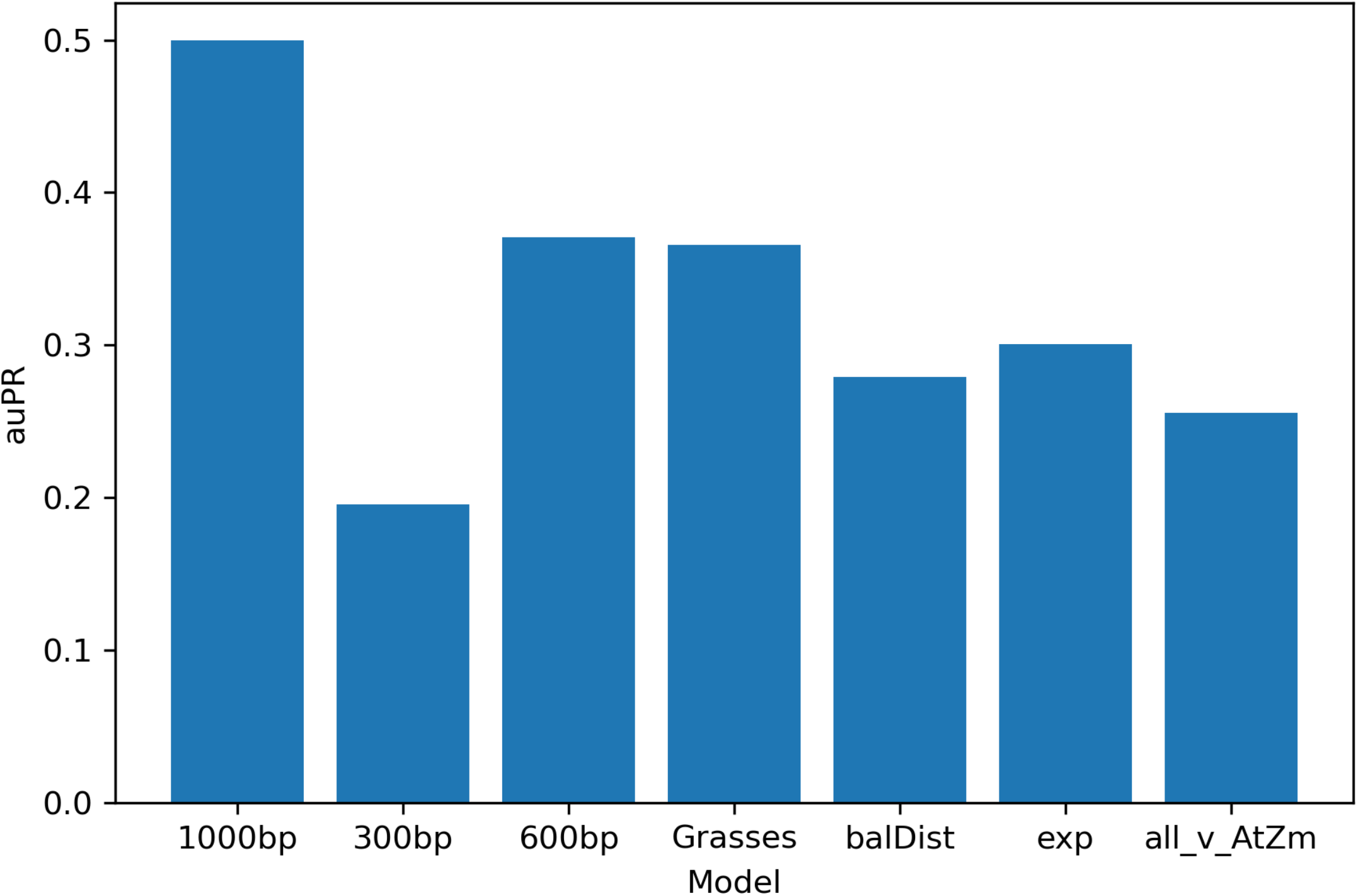
auPR of different across-species accessibility model configurations. From left to right: 1000bp windows, 300bp windows, 600bp windows, training and testing within only grasses, using a training set balanced on both accessibility and distance class, exponential activation on the convolutional layer, and testing on Arabidopsis and Maize while training on the rest.

**Figure S2:**
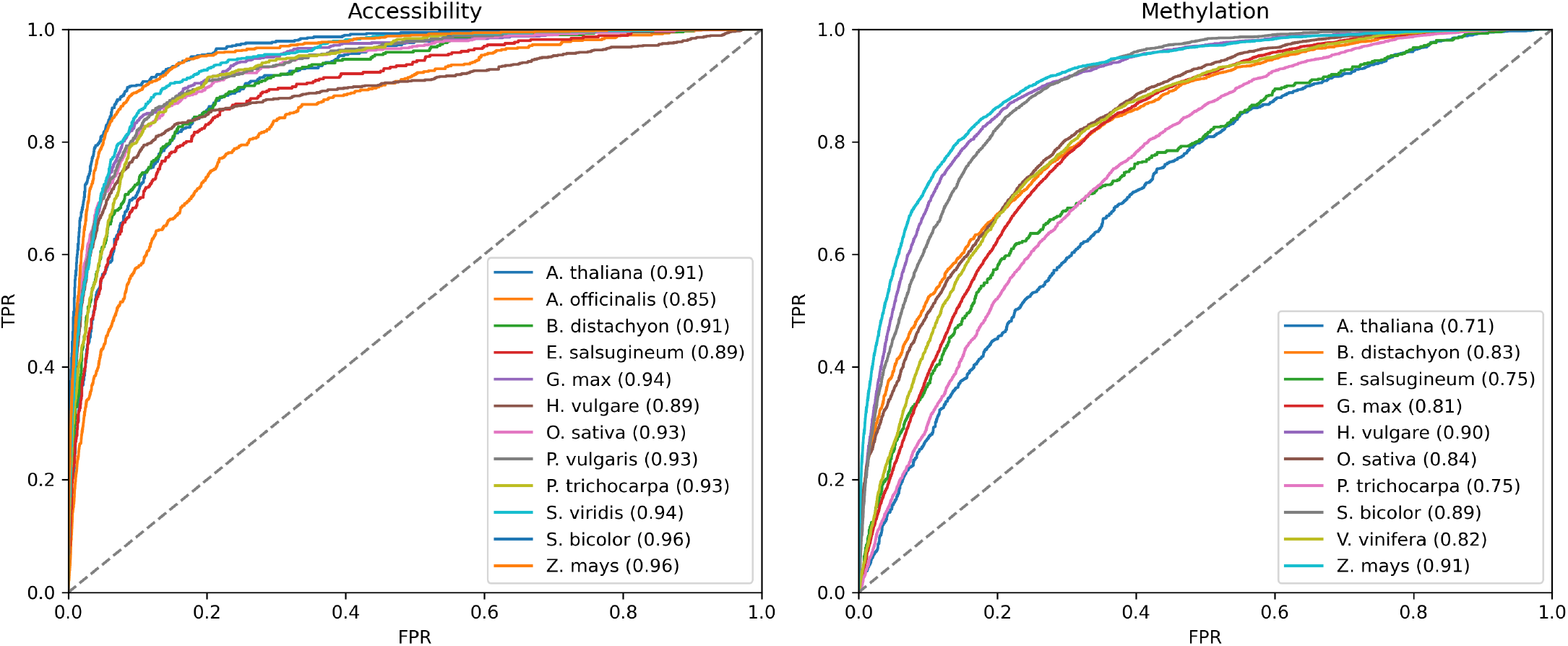
Receiver operating characteristic curves of the across-species models per hold-out species. The gray dashed line is the baseline expectation for a random classifier. The area under the curve is given inside the parentheses for each species in the legend.

**Figure S3:**
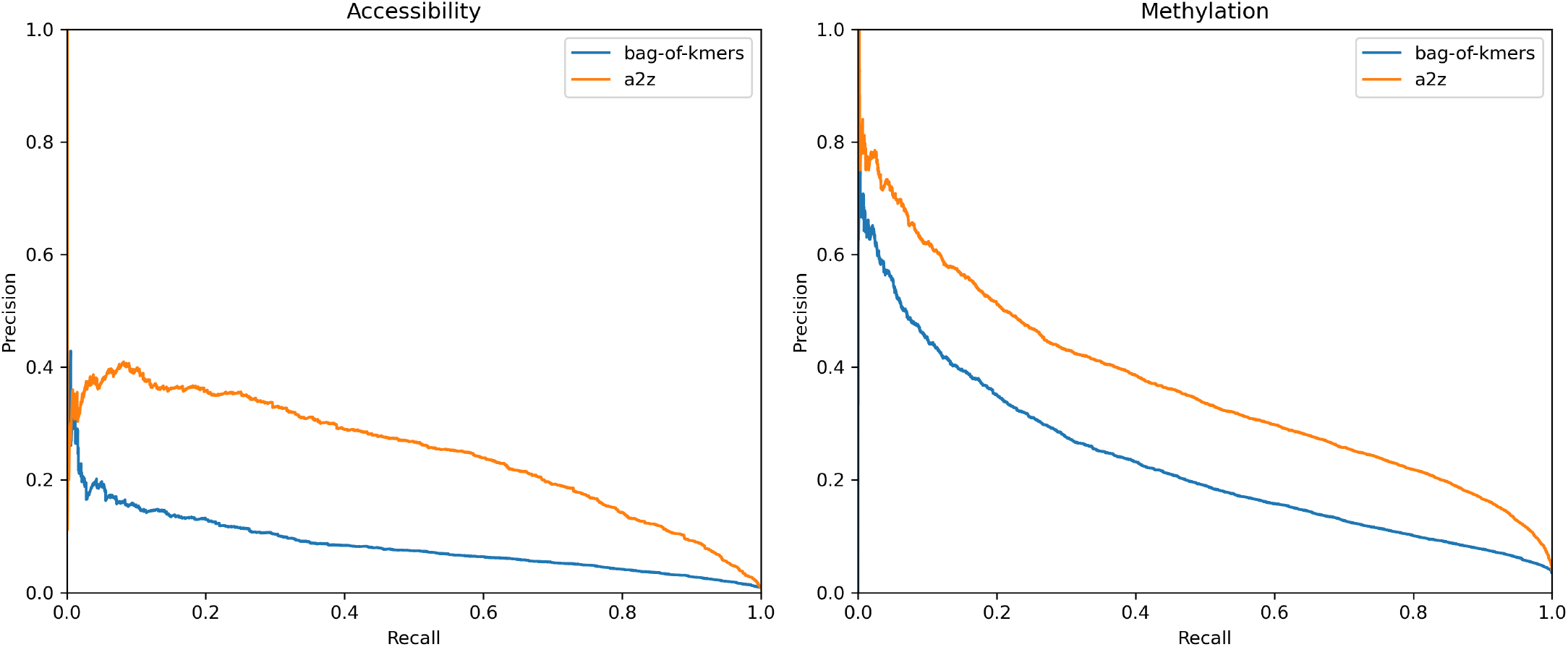
Precision-recall curve comparison of across-species a2z model with the bag-of-kmer model in *Z. mays*.

**Figure S4:**
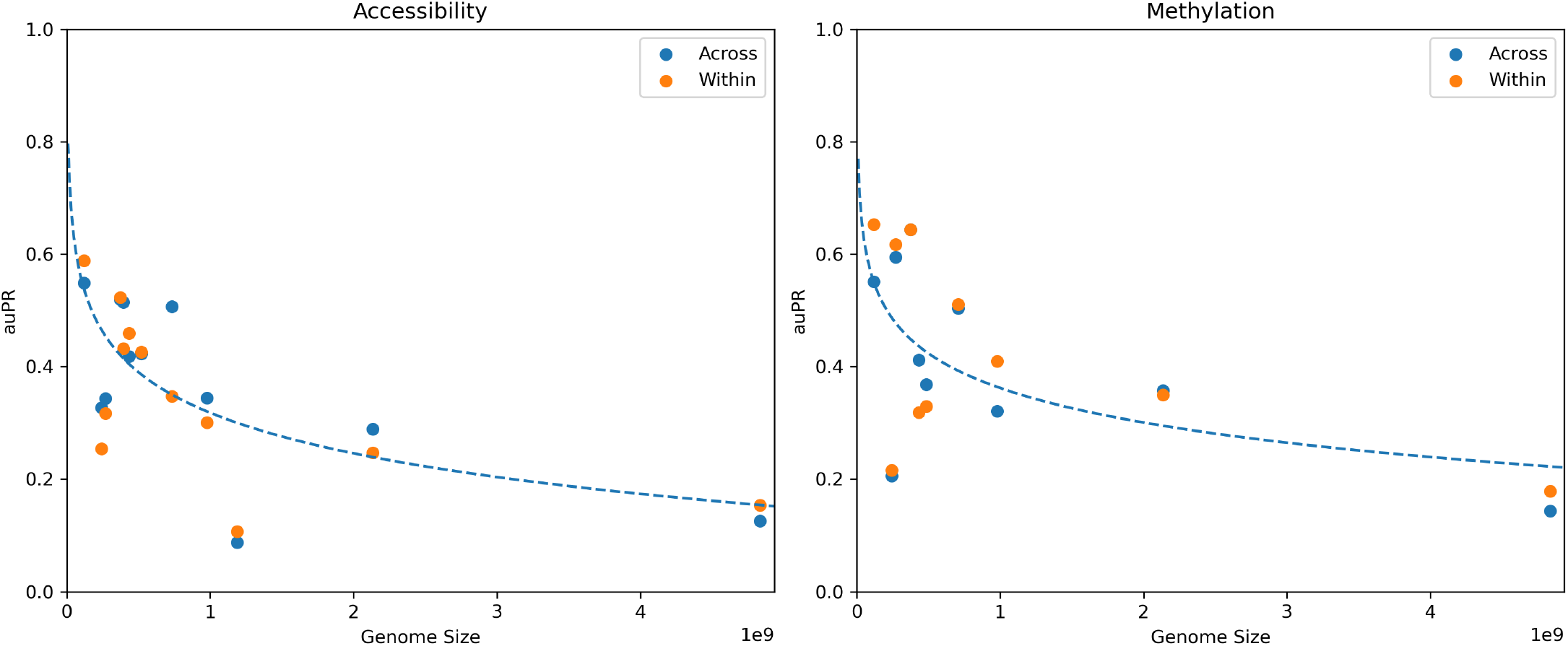
auPR of both model configurations with respect to genome size. The dashed line is an exponential fit to the across-species model auPR values.

**Figure S5:**
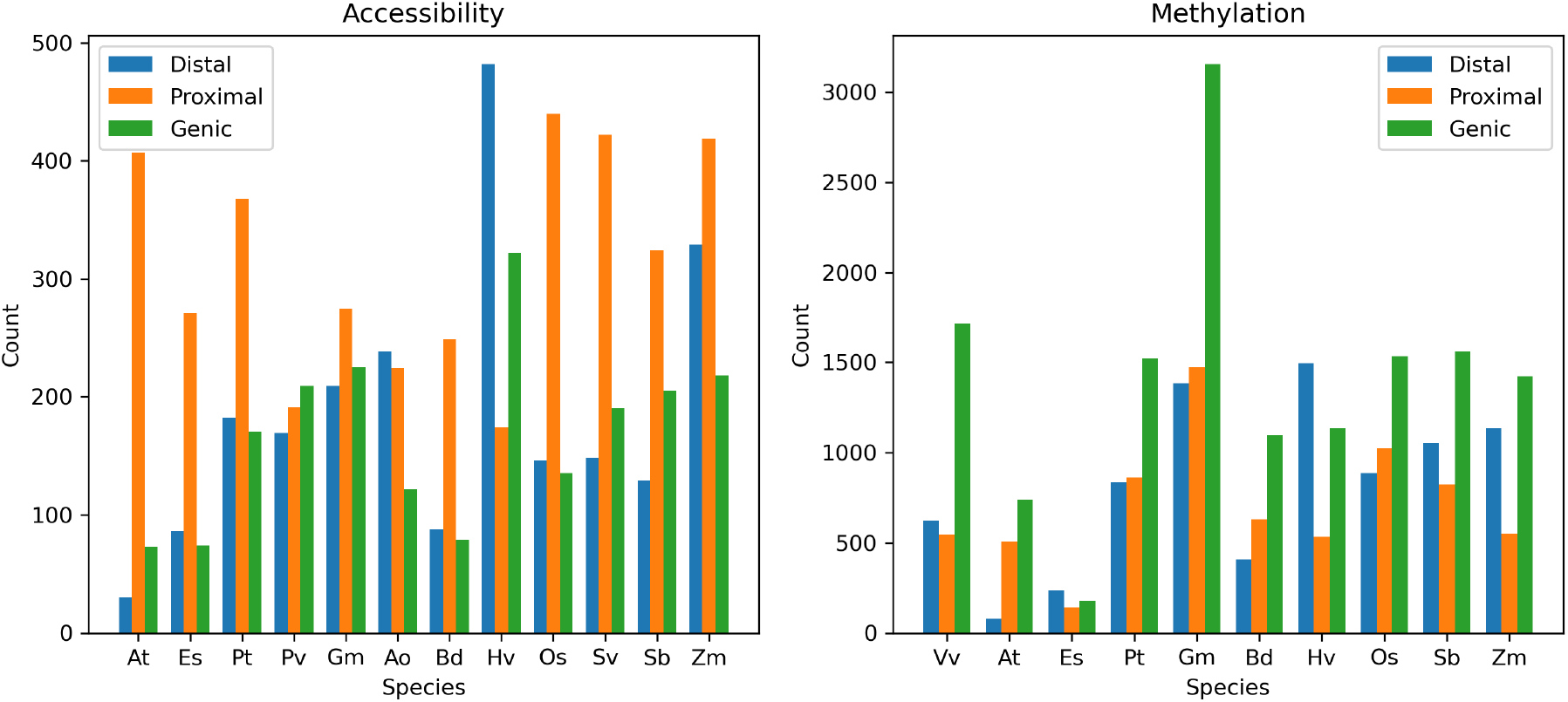
Counts of accessible regions in the across-species test sets by distance class and species.

**Figure S6:**
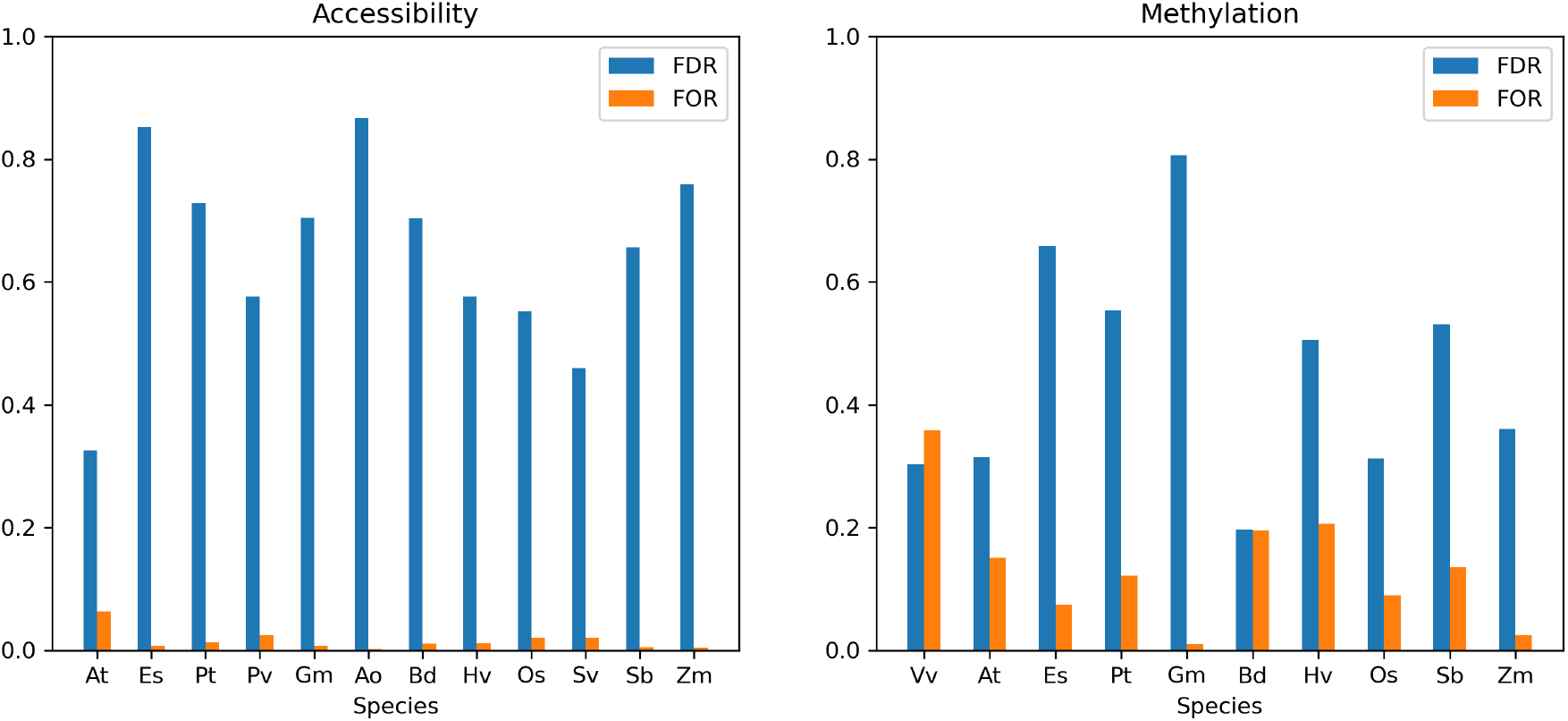
Comparison of false discovery rate (FDR) versus false omission rate (FOR) between models.

**Figure S7:**
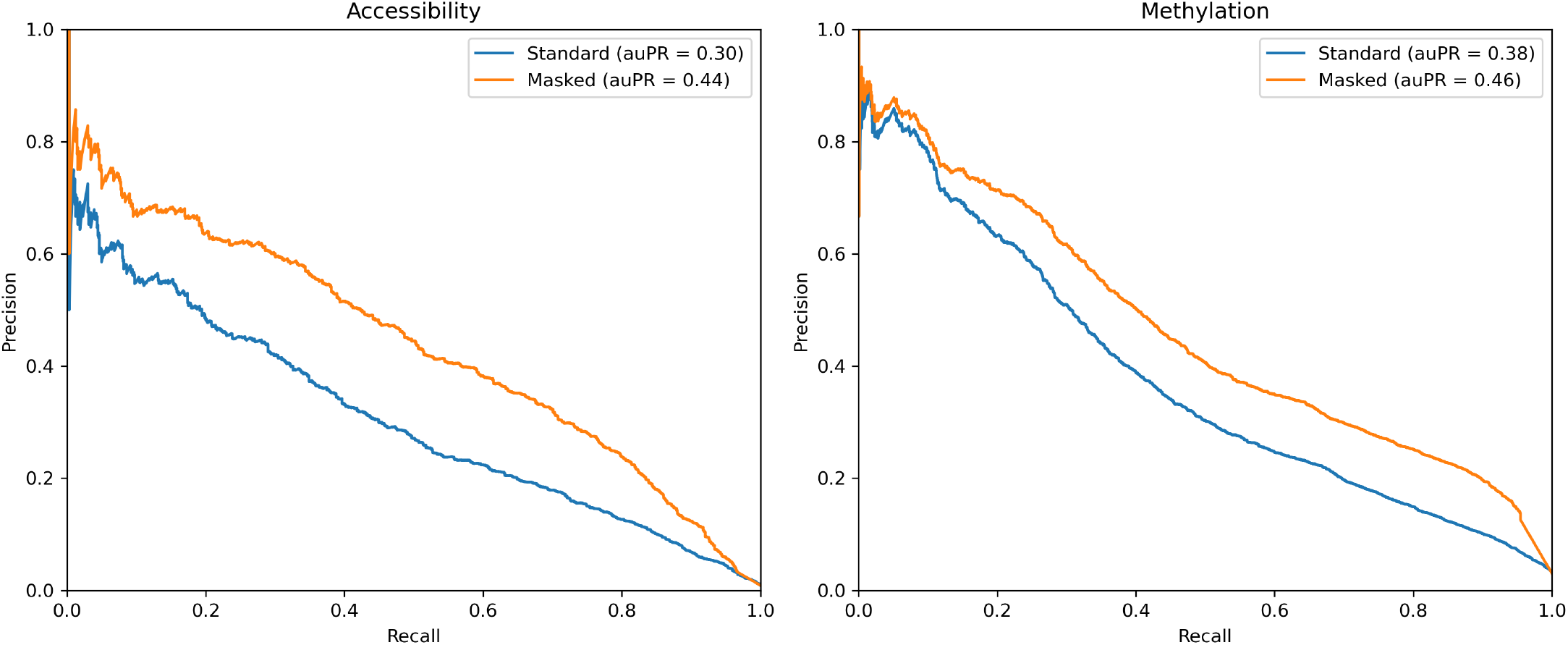
Precision-recall curve comparison between a2z model and the repeat-masked a2z model in *Z. mays*.

**Figure S8:**
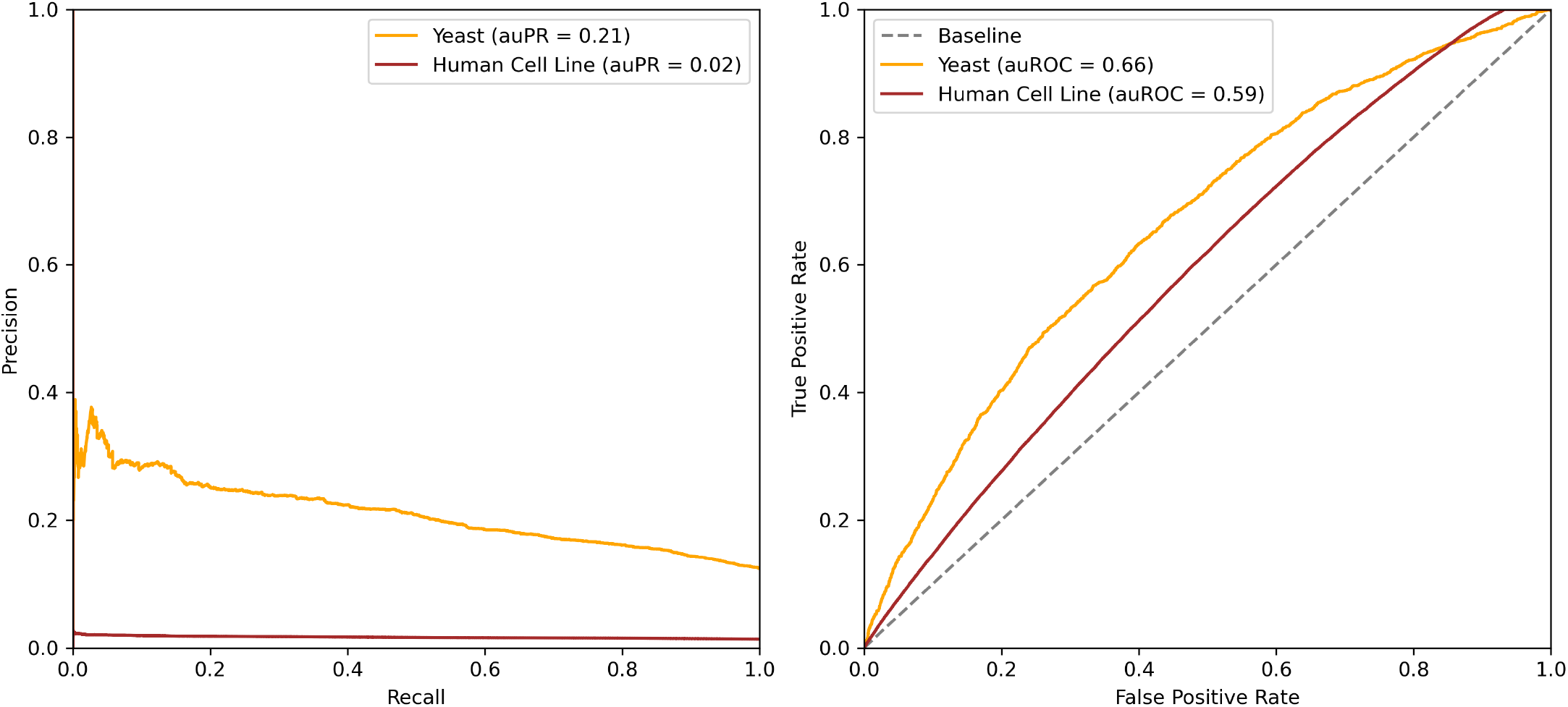
Precision-recall and receiver operating characteristic curves of an angiosperm-trained a2z model on yeast and a human cell line.

**Figure S9:**
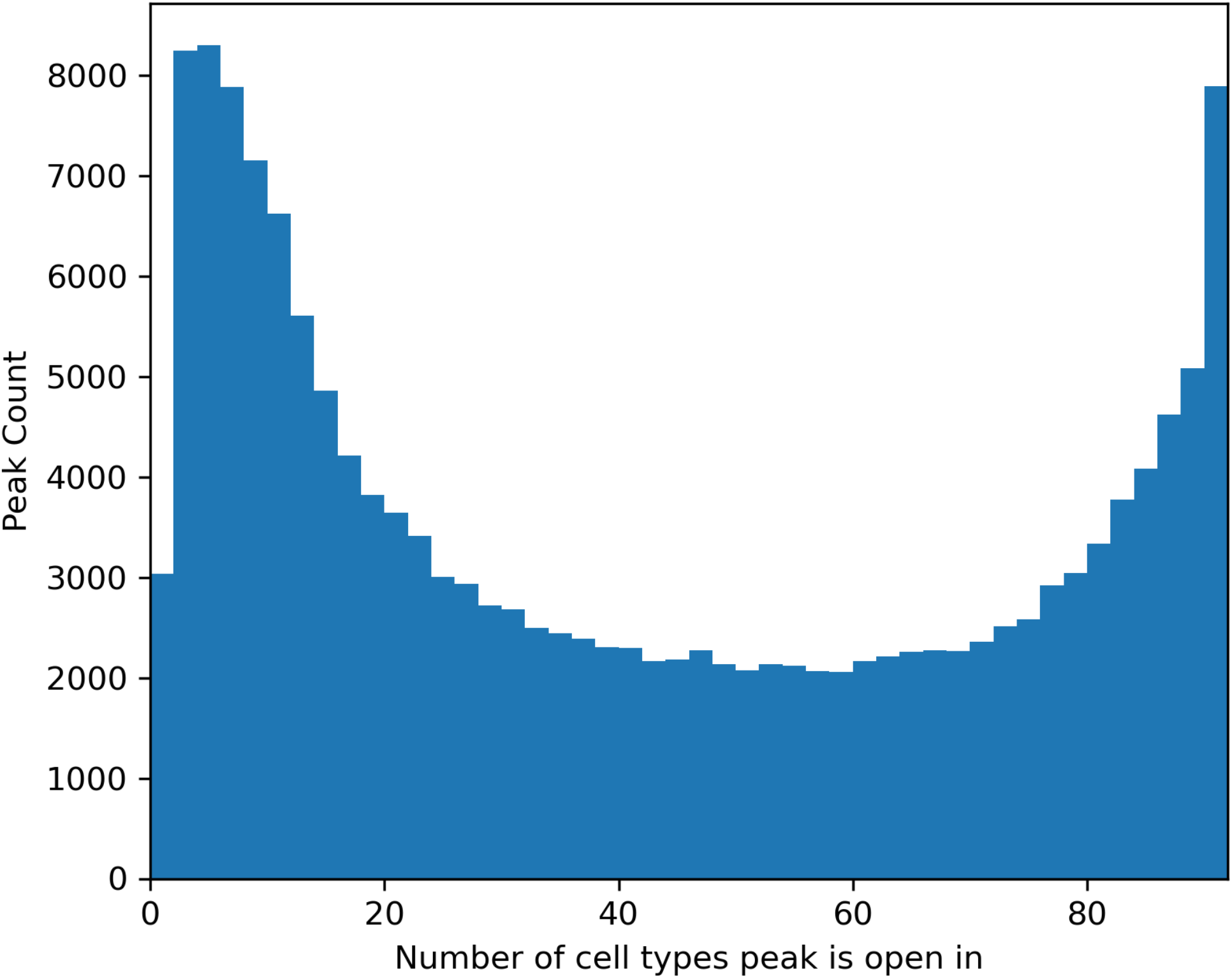
Cell type specificity of maize scATAC-Seq peaks.

**Figure S10:**
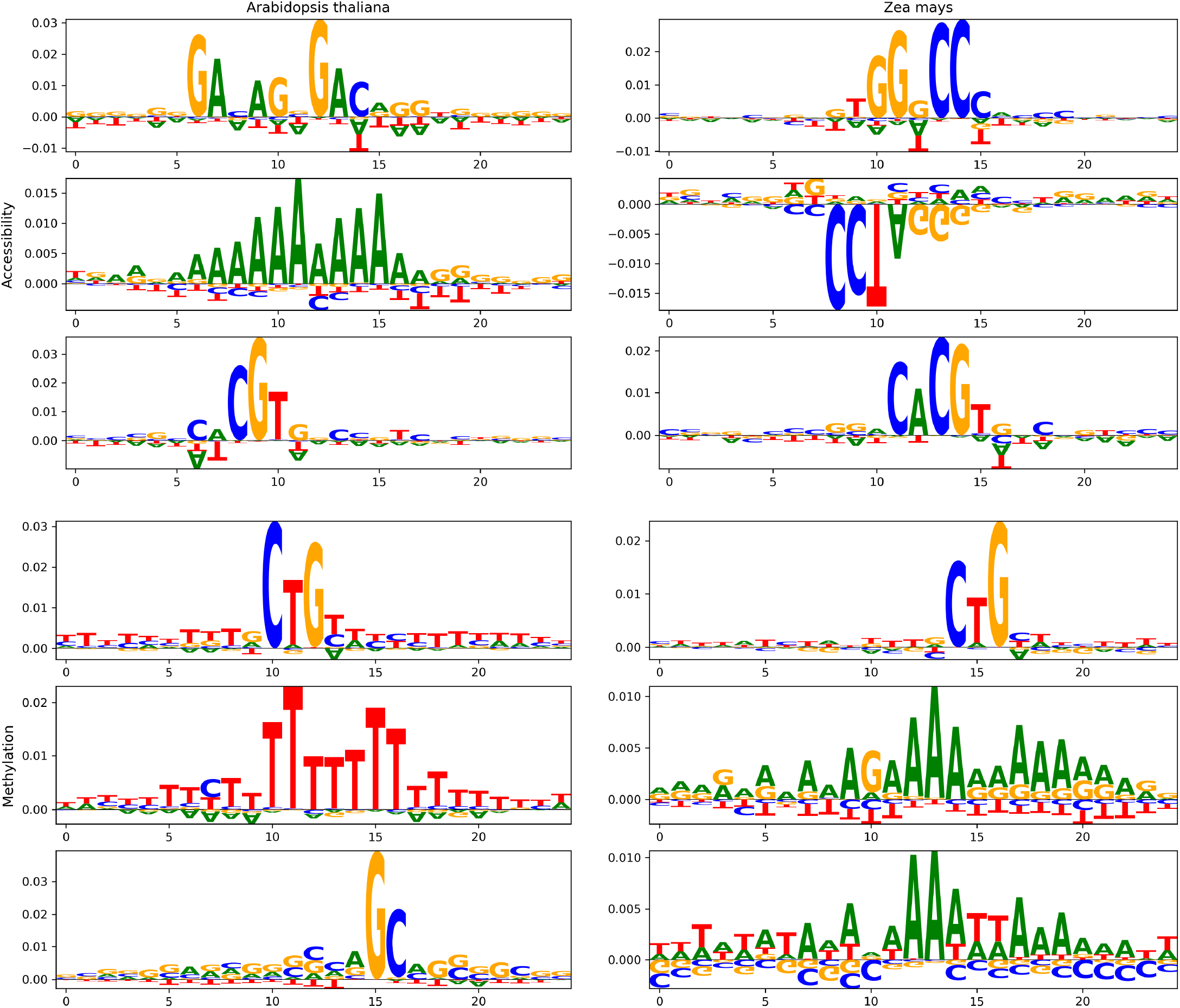
Top 3 TF-MoDISco patterns for four a2z models. *A. thaliana* is the left column, *Z. mays* is the right column. Accessibility is the top row and methylation is the bottom row. Within each species and chromatin feature combination the patterns are ranked from top to bottom by the number of supporting seqlets for that pattern.

**Figure S11:**
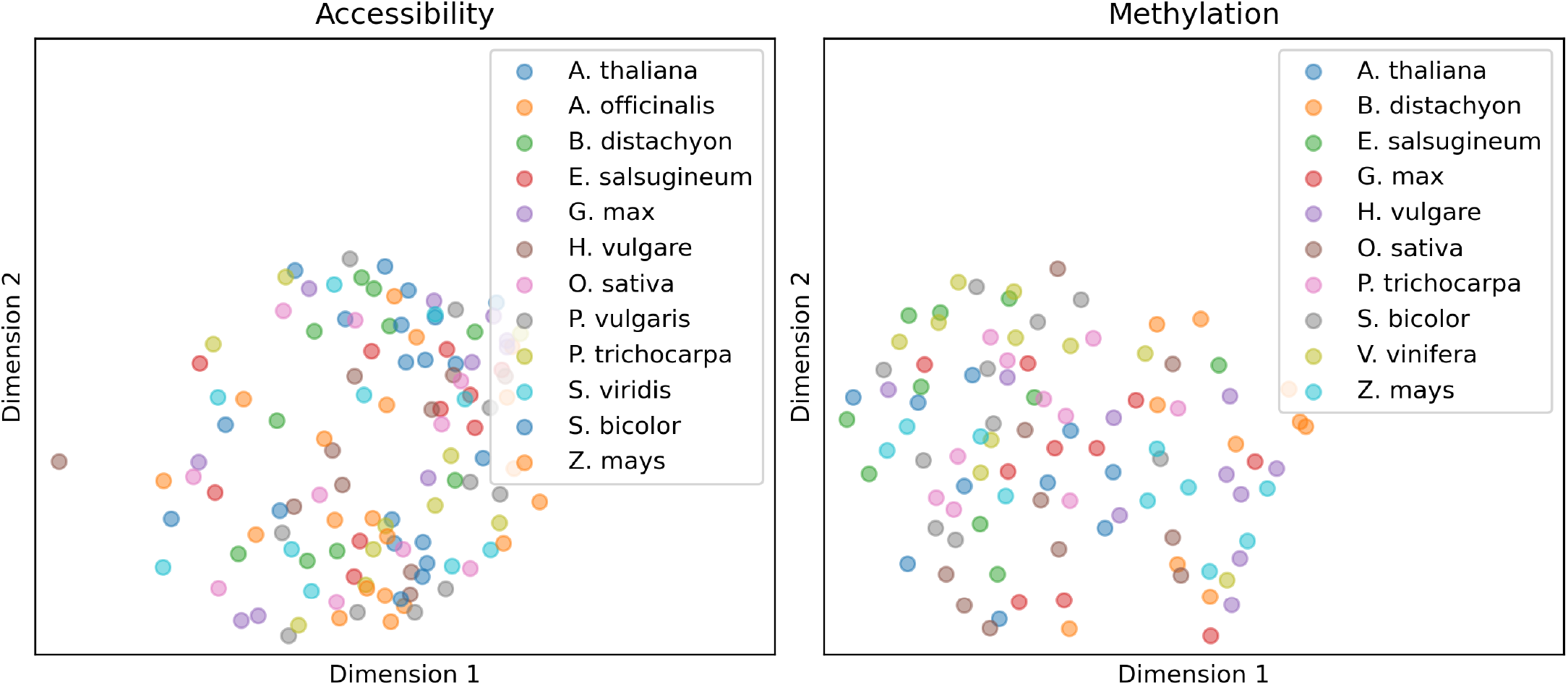
Multidimensional scaling of the top 10 high-effect medoid kmers for each species and chromatin feature model combination, colored by species.

**Figure S12:**
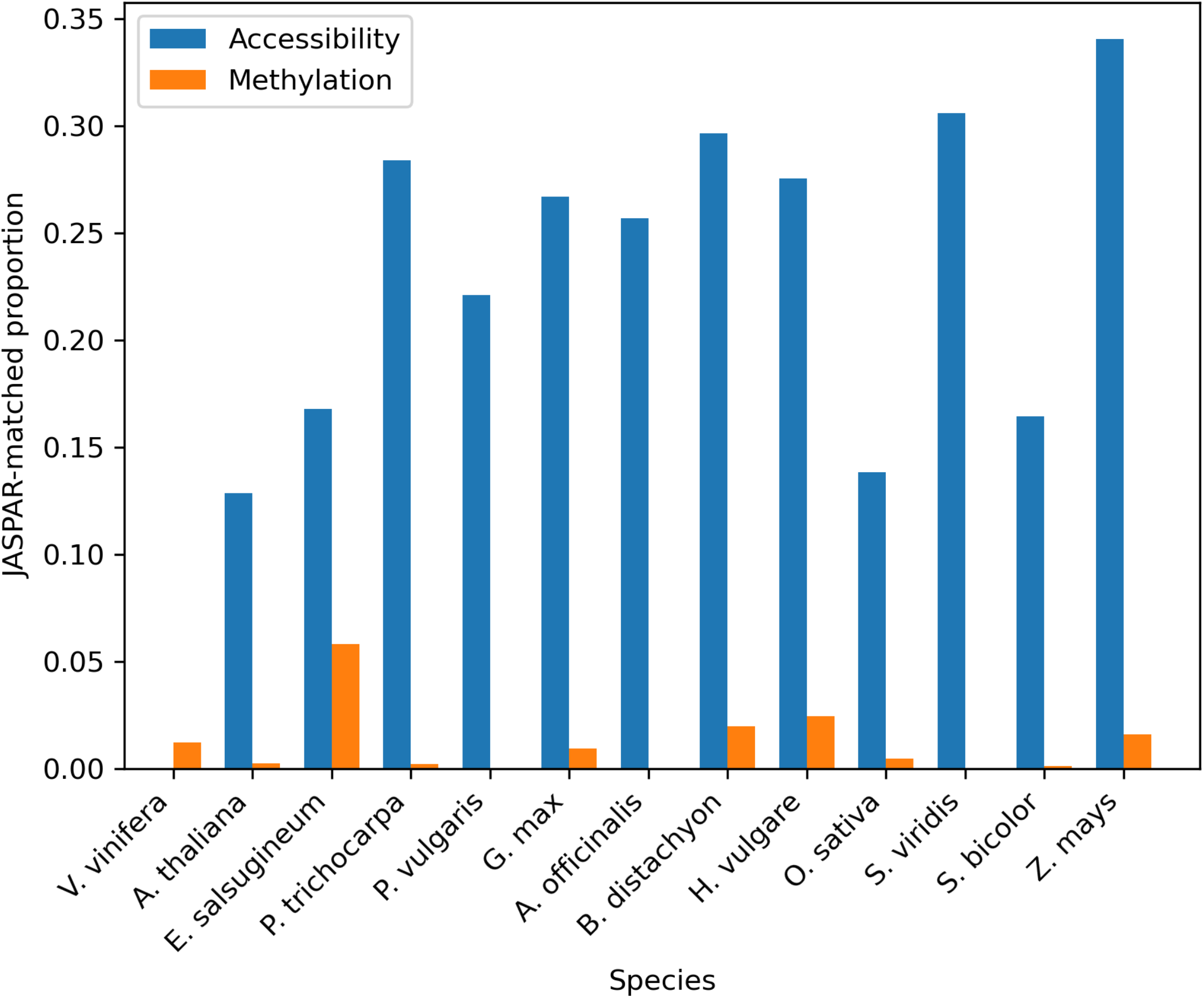
Proportion of high-effect kmers that significantly (q-value <= 0.05) matched JASPAR CORE *plantae* motifs with FIMO, grouped by species and chromatin feature.

## Notes

https://doi.org/10.5281/zenodo.5676313

### Competing Interest Statement

The authors have declared no competing interest.

